# Robust candidates for language development and evolution are significantly dysregulated in the blood of people with Williams syndrome

**DOI:** 10.1101/488155

**Authors:** Antonio Benítez-Burraco, Ryo Kimura

**Affiliations:** Department of Spanish, Linguistics, and Theory of Literature, Faculty of Philology, University of Seville, Seville, Spain; Department of Anatomy and Developmental Biology, Graduate School of Medicine, Kyoto University, Kyoto, Japan

**Keywords:** Williams syndrome, blood transcriptional profile, language disorders, language evolution

## Abstract

Williams syndrome (WS) is a clinical condition entailing cognitive deficits and with an uneven language profile, which has been object of intense inquiry over the last decades. Although WS results from the hemideletion of around two dozens of genes in chromosome 7, no gene has been yet probed to account for, or contribute significantly to, the language problems exhibited by the affected people. In this paper we show that robust candidates for language disorder and for language evolution in the species, located outside the hemideleted region, are up– or downregulated in the blood of subjects with WS. Most of these genes play a role in the development and function of brain areas involved in language processing, which exhibit structural and functional anomalies in people with the condition. Overall, these genes emerge as robust candidates for language dysfunction in WS.

## Introduction

Williams syndrome (WS) is a clinical condition resulting from a hemizygous deletion of 1.5 to 1.8 Mb on 7q11.23, which affects to nearly 30 genes (Korenberg et al., 2000; Pober, 2010). The affected people exhibit a distinctive behavioral and cognitive profile, with enhanced sociability, mental retardation, impaired spatial cognition, and spared musical abilities (Reilly et al., 1990; Udwin and Yule, 1991; Bellugi et. al., 1999; Galaburda et al., 2002; Levitin et al., 2005). Language abilities are significantly preserved in people with WS compared to other neurodevelopmental disorders, although they are delayed or impaired across different levels compared to the neurotypical population (Karmiloff-Smith and Mills, 2006; Brock, 2007; Mervis and Becerra, 2007; Martens et al., 2008 for good reviews). Children with WS experience problems mostly with irregular forms and complex syntax; conversely, they usually excel on expressive vocabulary (including semantic organization and fluency), although they have problems with word definitions (Volterra et al., 1996, Mervis et al., 1999, Purser et al., 2011; Van Den Heuvel et al., 2016; see Mervis and Becerra 2007 for discussion). Nonetheless, as with other aspects of the cognitive profile of this condition, no robust gene-to-phenotype associations have been established in the language domain. To date, the most promising candidates for language dysfunction in WS are *GTF2I, BAZ1B*, and *LIMK1*. In particular, *GTF2I*, which encodes a regulator of transcription, has been repeatedly related to the behavioral and cognitive disabilities that are typically found in this condition and that have an impact on language function (Morris et al., 2003; Tassabehji et al., 2005; Sakura et al., 2011; Hoeft et al., 2014). The gene has been associated to autism spectrum disorder (ASD) too (Malenfant et al. 2012). Its adjacent paralog, *GTF2IRD1*, has been related to altered vocalizations, among other features (Howard et al., 2012). Interestingly too, *BAZ1B* happloinsuficiency explains almost 50% of transcriptional dysregulation in WS neurons, with BAZ1B target genes being enriched in functions related to neurogenesis and neuron differentiation (Lalli et al., 2016). Regarding *LIMK1*, it regulates synaptic plasticity and long-term memory (Todorovski et al. 2015), and its hemideletion has been hypothesized to account for the deficits in spatial cognition in combination with other genes (Gray et al., 2006; Smith et al., 2009). Still, these links are quite vague, if one considers our remarkable understanding of the genetic underpinnings of human language, language disorders, and language evolution (see Scharff and White, 2004; Li and Barlett, 2012; Benítez-Burraco, 2013; Graham et al., 2015; Fisher, 2017; Murphy and Benítez-Burraco, 2017; 2018 for reviews). Examining how these robust candidates for language dysfunction and evolution behave in WS should help refine our view of the molecular causes of language dysfunction in this condition. One general reason is the deep link that exists between evolution and (abnormal) development, in the spirit of evo-devo theories. One particular reason is that although in this condition the number of hemideleted genes is quite reduced, changes in the dosage of hundreds or even thousands of other genes can be expected, with a potential impact on language abilities. Recently Kimura and colleagues (2018) confirmed that the dysregulation of several co-expression modules involving dozens of genes outside of the 7q11.23 region seemingly accounts for the complex phenotypes observed in WS patients. Importantly, they found *BCL11A*, a gene associated with speech disorders, among the hub genes in the top WS-related modules.

In this paper we conduct a more focused research on the potential dysregulation of genes related to language outside the WS region as a possible explanation of the distinctive language profile of the affected people. We found that significant differences exist in the blood of subjects with WS compared to neurotypical controls in the expression levels of robust candidates for language evolution and development.

## Method

The list of core candidates for language (abnormal) development and language evolution (Supplemental table 1) encompasses two subsets of genes. The first subset consists of strong candidates for language disorders, in particular, dyslexia and specific language disorder (SLI), as listed by Paracchini and colleagues (2016), Pettigrew and colleagues (2016) and Chen and colleagues (2017). The second subset consists on strong candidates for language evolution, as compiled by Boeckx and Benítez-Burraco (2014a; 2014b) and Benítez-Burraco and Boeckx (2015). These are genes involved in the globularization of the human skull/brain and the cognitive changes accounting for our species-specific ability to learn and use languages (aka our *language-readiness*), and fulfil the following criteria: they have changed (and/or interact with genes that have changed) after our split from Neanderthals/Denisovans (including changes in their coding regions and/or their epigenetic profile); they play some known role in brain development, regionalization, wiring, and/or function; and/or they are candidates for language dysfunction in broad cognitive disorders, particularly, ASD and schizophrenia (SZ) (see Benítez-Burraco and Murphy, 2016; Murphy and Benítez-Burraco, 2016; 2017 for details about their role in language processing).

**Table 1.** Genes that are significantly dysregulated in the blood of subjects with WS (FDR <0.05, |FC| > 1.2).

| Gene name | logFC | FDR Pvalue |
| --- | --- | --- |
| *PVALB* | -0,651 | 1,65E-03 |
| *DOCK4* | -0,500 | 2,00E-03 |
| *NFXL1* | -0,447 | 7,77E-03 |
| *ZNF385D* | -0,424 | 6,59E-06 |
| *DUSP1* | -0,380 | 6,66E-03 |
| *CEP63* | -0,370 | 3,43E-05 |
| *PAX3* | -0,348 | 4,13E-02 |
| *ITGB4* | -0,298 | 7,37E-03 |
| *PLAUR* | -0,295 | 4,88E-02 |
| *FGFR1* | 0,286 | 7,44E-03 |
| *SETBP1* | 0,299 | 1,93E-03 |
| *SIX3* | 0,310 | 1,55E-03 |

The gene expression profiling data of peripheral blood in patients with WS was obtained from Gene Expression Omnibus (GSE 89594). The Benjamini-Hochberg method (Benjamini and Hochberg, 1995) was used to calculate the false discovery rate (FDR). Genes were considered to be differentially expressed genes (DEG) when the FDR <0.05 and the |fold change (FC)| > 1.2. Statistical overrepresentation was evaluated using Fisher’s exact test.

## Results

We found that candidates for language (abnormal) development and language evolution are significantly dysregulated in the blood of subjects with WS (p=1.1e-7 by Fisher’s exact test). Table 1 shows the genes that are significantly up– or down-regulated compared to controls (FDR <0.05, |FC| > 1.2).

In order to check the specificity of this set of genes in relation to language we conducted a functional enrichment analysis with Enrichr (amp.pharm.mssm.edu/Enrichr; Chen et al., 2013; Kuleshov et al., 2016). Our results (Supplemental table 2) show that the dysregulated genes mainly contribute to the cytoskeleton activity and are significantly involved in cell proliferation and migration, including neuroblast proliferation. Regarding their molecular function, they typically participate in protein modification, particularly via (tyrosine) kinase phosphatase and (tyrosine) kinase binding activities, but also in gene regulation, via transcription cofactor binding. Interestingly, these genes are significantly associated to aberrant processes impacting on brain development, like abnormal neural tube morphology and neural crest cell migration, as well as decreased forebrain size and abnormal midbrain development. Likewise, they are associated to clinical symptoms mostly impacting on craniofacial morphology, like malar flattening, midface retrusion, shallow orbits, or depressed nasal bridge. Finally, these genes are predicted to be preferentially expressed in the ectoderm, the cranium, the retina, and the neural crest. According to the Human Brain Transcriptome Database (http://hbatlas.org) all these genes are expressed in the brain, particularly in the thalamus and the cerebellum (see Supplemental table 3). The thalamus functions as a sort of relay center to connect many brain areas involved in language processing (Wahl et al., 2008; Murdoch, 2010; David et al., 2011) and changes in the thalamus have been claimed to contribute to the evolutionary emergence of our language-ready brain (see Boeckx and Benítez-Burraco, 214a for details). Similarly, the cerebellum plays a key role in language processing and is impaired in language-related pathologies (Vias and Dick, 2017; Mariën and Borgatti 2018). People with WS exhibit cerebellar volume alterations that are seemingly associated with their cognitive, affective and motor distinctive features (Osório et al., 2014). In the same vein, the thalamus exhibits structural and functional differences with the neurotypical population, including disproportionately reduced volumes and decreased gray matter (Chiang et al., 2007; Campbell et al., 2009) and enhanced thalamic activity (Mobbs et al., 2007; Bódizs et al., 2012).

Nearly one third of the genes found downregulated in the blood of subjects with WS are candidates for dyslexia (*DOCK4, ZNF385D, CEP63*) and/or for SLI (*DOCK4, NFXL1*). As other members of the Dock family, DOCK4 regulates cytoskeleton assembly, and cell adhesion and migration (Gadea and Blangy, 2014). Specifically, DOCK4 has been shown to be involved in neuronal migration and neurite differentiation (Ueda et al., 2008; Xiao et al., 2013), via interaction with the actin-binding protein cortactin (Ueda et al., 2013). Knockdown of *Dock4* in mice abolishes commissural axon attraction by Shh (Makihara et al., 2018). The gene has been related to neuronal migration and neurite outgrowth anomalies linked to developmental dyslexia (Shao et al. 2016), although it is also associated with ASD (Pagnamenta et al., 2010) and SZ (Alkelai et al., 2012). GWAs have associated markers in *ZNF385D* to the co-occurrence of reading disability and language impairment (Eicher et al., 2013), but also to negative symptoms in SZ (Xu et al., 2013). *CEP63* is required for normal spindle assembly, being involved in maintaining centriole number and establishing the order of events in centriole formation (Brown et al., 2013). Besides its association with dyslexia (Einarsdottir et al., 2015), the gene is also a candidate for primary microcephaly (Marjanović et al., 2015), a feature that is commonly found in subjects with WS (Jernigan and Bellugi, 1990; Schmitt et al., 2001; Thompson et al., 2005; Jackowski et al., 2009). Finally, variants of *NFXL1*, which is predicted to encode a transcription factor, confer a risk for SLI (Villanueva et al., 2015). The gene is highly expressed in the cerebellum (Nudel, 2016).

Regarding the candidates for language evolution that we have found downregulated in the blood of subjects with WS, *DUSP1* is involved in vocal learning in songbirds (Horita et al. 2010, Horita et al. 2012). *PVALB* encodes a calcium-binding protein that is structurally and functionally similar to calmodulin and that is involved in hippocampal plasticity, learning and memory (Donato et al., 2013). Interestingly enough, the inactivation of Pvalb-expressing interneurons in the auditory cortex alters response to sound, strengthening forward suppression and altering its frequency dependence (Phillips et al., 2017). Inhibition of PVALB-expressing GABAergic interneurons results in complex behavioral changes related to the behavioral phenotype observed in SZ (Brown et al., 2015). Importantly, some of the key changes that contributed to the emergence of our language-readiness involved GABAergic signaling (discussed in detail in Boeckx and Benítez-Burraco, 2014a), which are vital for oscillatory processes underlying language processing (Bae et al., 2010; see Murphy and Benítez-Burraco, 2018 for details). Reduction in *PVALB* expression in interneurons has been also found in mouse models of ASD (Filice et al., 2016), specifically, in the Cntnap2-/-model (Lauber et al., 2018). *CNTNAP2* is a direct target of FOXP2, the renowned “language gene” (Vernes et al., 2008; Adam et al., 2017) and regulates language development in non-pathological populations too (Whitehouse et al. 2011, Whalley et al. 2011, Kos et al. 2012). Also mice lacking *Plaur* have significantly fewer neocortical parvalbumin-containing GABAergic interneurons, this being associated with impaired social interactions (Bruneau and Szepetowski, 2011). *PLAUR* is a target of FOXP2 too (Roll et al. 2010), but also an effector of SRPX2, another of FOXP2’s targets (Royer-Zemmour et al. 2008) and a candidate for speech dyspraxia (Roll et al. 2006). Concerning PAX3, this gene is expressed in the neural crest and is a candidate for Waardenburg syndrome, a clinical condition entailing sensorineural hearing loss and developmental delay (Tassabehji et al., 1992; Chen et al., 2010). Finally, ITGB4 encodes the integrin beta 4 subunit, a receptor for the laminins, including FLNA (Travis et al, 2004), an actin-binding protein needed for cytoskeleton remodeling and neuronal migration (Fox et al. 1998), which in turn binds one of the proteins bearing fixed changes in humans compared to extinct hominins (Pääbo, 2014, table S1).

Lastly, among the genes shown to be upregulated in the blood on WS subjects, we found the SLI candidate *SETBP1*, as well as *FGFR1* and *SIX3*. SETBP1 is also a candidate for Schinzel-Giedion syndrome, a clinical condition entailing occasional epilepsy and severe developmental delay (Ko et al. 2013, Miyake et al. 2015). Mutations on this gene have been also associated to behavioral and social deficits (Coe et al. 2014). The Integrative Nuclear FGFR1 Signalling (INFS) has been hypothesized to be one of the neurodevelopmental pathways on which multiple SZ candidates converge, regulating numerous neurotransmitter systems and neural circuits (Stachowiak et al., 2013). Finally, *SIX3* contributes to regulate the relative size of the telencephalon versus the thalamus (Lavado et al. 2008, Sylvester et al. 2010). Interestingly, Six3 regulates Shh (Jeong et al., 2008), one robust candidate for microcephaly that has been positively selected in the human lineage (Dorus et al. 2004), but also interacts with several genes relevant for our language-ready brain (Benítez-Burraco and Boeckx, 2015)

## Discussion

Deciphering the exact molecular causes of language dysfunction in WS is still pending. At present, none of the genes hemideleted in this condition has been demonstrated to play a central role in language processing. At the same time, hundreds of genes outside the WS critical region have been found to be dysregulated in subjects with this condition. In this paper we have shown that these genes are significantly enriched in core candidates for language disorders and language evolution, which emerge as robust candidates for language dysfunction in WS.

## Supporting information

## Acknowledgements

This research was funded by the Spanish Ministry of Economy and Competitiveness (grant FFI2016-78034-C2-2-P [AEI/FEDER, UE] to ABB)

